# The unusual gene order in the Echinoderm Hox cluster is related to the embryo and larva symmetries

**DOI:** 10.1101/016923

**Authors:** Spyros Papageorgiou

## Abstract

**Background:** Hox gene collinearity relates the sequential location of Hox genes in the 3’ to 5’ direction on the chromosome with the linear arrangement of the body elements along the anterior-posterior (A/P) axis of bilaterian embryos. This spatial Hox gene collinearity has been almost universally respected in diverse organisms like worms, insects or vertebrates. It is therefore surprising that the above well established collinearity rule is violated in the case of Echinoderms. No explanation of this violation is apparent. Here a hypothesis is put forward which provides a cue to understand the abnormal serial gene location in the sea urchin disorganized Hox cluster.

**Results:** Bilateral symmetry along the A/P embryo axis is established at the very early stages of ontogeny of the sea urchin. For the subsequent developmental stages, rotational symmetry emerges in the vestibula larva. In analogy to the linear A/P case, the circular topology of modules might be a reflection of the architectural restructuring of the Hox loci where the 3’ and 5’ ends of the Hox cluster approach each other so that a closed contour of the chromatin fiber is formed. At a later stage, the break and opening of the cluster contour at the level of Hox4 combined with the rotational symmetry leads to the observed Hox gene sequence that violates the standard 3’ to 5’ collinearity.

**Conclusion:** The unusual gene series manifests the congruence of Hox gene sequence in the cluster with the circular arrangement of the sea urchin primary podia. Accordingly, the Hox sequence after the break at Hox4 is not a violation but an extension of Hox gene collinearity to animals with rotational symmetry.

## Introduction

Hox genes play an important role in the normal development of all bilaterian animals humans included. In particular the organization of Hox genes in clusters led to the surprising observation of **Hox gene collinearity**: the genes are numbered along the 3’ to 5’ direction on the chromosome and their expressions follow the same order in the anterior-posterior (A/P) direction of the embryo (**spatial collinearity**) [1]. This property is particularly evident in organized and compact clusters as in the case of vertebrates [2]. In this case it is further established that the Hox genes obey **temporal collinearity**, where gene counting starts from the 3’ end on the chromosome: the first gene in the sequence (Hox1) is expressed first, followed by Hox2, then by Hox3 etc.

The explanation of collinearity has been at the center of interest of specialists on Development and Evolution. The last fifteen years or so genetic engineering techniques were developed (particularly by D. Duboule and coworkers, see e.g.[3,4]) enabling the deletion or duplication of Hox genes in the cluster of vertebrates. These experiments shed light to some facets of the mechanism responsible for the collinearity of vertebrate Hox genes. An explanatory ‘two-waves model’ has been formulated by Duboule and coworkers [3-6]. It is based on well established biomolecular processes according to which enhancers located in a telomeric ‘landscape’ activate the Hox genes of the cluster [5]. Besides the telomeric landscape the cluster is flanked by a centromeric landscape which is acting negatively and balances the influence from the telomeric side [4,5].

A different approach is proposed by a ‘biophysical model’. According to this model, physical forces are generated which translocate the Hox genes one after the other from a secluded location where the genes are inactive to an area where transcription is possible [7-9].

The collinearity rules apply successfully along the anterior-posterior axis of the bilaterian animals at their early developmental stages. In this case collinearity relates two structures of distinct scales: on one hand, at the chromosome subcellular level, the genes are deployed on a line with distinct ends: the telomeric (3’) and the centromeric (5’) ends of the Hox cluster. At the other hand, the multicellular body units develop along the anterior-posterior axis of the embryo. This coordinated interrelation indicates that somehow the two multiscale structures communicate and affect each other [8-10].

## Ontogeny of Echinoderms

After the completion of the sea urchin sequencing in 2006 by a specific Consortium, the study of Echinoderms and in particular the sea urchin Hox gene cluster is advancing quite rapidly [11]. A striking result from this DNA analysis is the unusual gene order in the sea urchin Hox cluster as compared to the vertebrate Hox cluster [12]. The gene locations in these two characteristic clusters are depicted in Fig.1. In the unusual sea urchin cluster, Hox1 to Hox3 have been translocated to the 5’ end while the other genes are inverted and translocated to the 3’ end as shown in Fig.1b [13].

**Figure 1:**
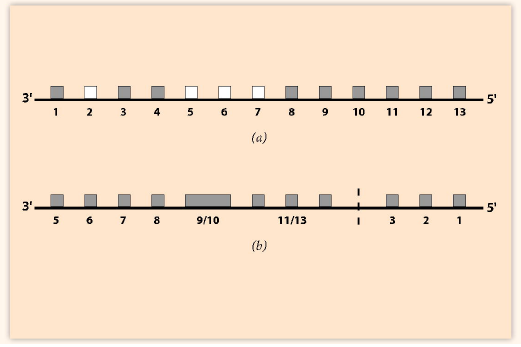
Schematic gene order in the Hox clusters. The real distances between the genes (pink squares) are not equal. Some genes are missing (open squares). (a) A typical normal cluster – the mouse HoxD cluster. Hoxd2, Hoxd5, Hoxd6, Hoxd7 are missing. (b) The unusual *Sea urchin* Hox cluster. Hox4 is missing (dashed line) [13].

In normal collinearity the ontogenic anterior-posterior (A/P) axis of the embryo is paralleled to the 3’ to 5’ axis of the Hox cluster. According to an analysis of the Hox loci of different species, the vertebrates possess the more compact and well organized clusters [2]. In contrast, the Hox clusters of the Echinoderms are spread in a much longer area of the genome and their structure appears disorganized [2].

In Echinoderms, bilateral symmetry and anterior-posterior patterning clearly appear at the very early stages of their ontogeny [13,14]. Subsequently and for the sea urchin larva stages, rotational and radial symmetries emerge [14]. The larva circular organization is superimposed on the A/P axis and the embryonic anterior and posterior ends cannot be distinguished in the new modules on the circular pattern [14] (Fig.2a). In this case the usual Hox gene collinearity breaks down [12,13].

**Figure 2:**
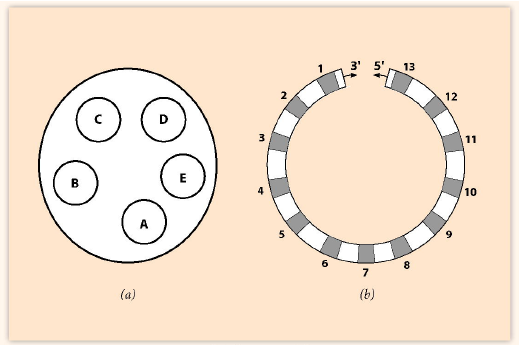
(a) Schematic representation of the oral view of the *H. purpurescens* vestibula larva (44h) with the 5 podia A, B, C, D, E [14]. (b) Circular arrangement of the gene fiber with the 13 Hox genes locations (see Fig.1a).

It is tempting to presume that the emerging symmetries of the sea urchin larvae are related to the violation of the usual spatial collinearity rule. How this interrelation is achieved remains to be clarified. The lengthy and disorganized structure of the sea urchin Hox cluster might also contribute to this violation [2,13]. In the present short work I propose a mechanism leading to the unusual gene order of the sea urchin Hox cluster.

In recent years the architectural restructuring of the Hox loci has been intensively studied [5,6]. It has been noticed that strong reorganization of the cluster occurs at the different stages of Hox activation. In this spirit it is assumed that the initial arrangement of the Hox genes on a straight line, as shown in Fig.1a, can be restructured to a closed contour where the 3’ and 5’ ends of the cluster come close to each other (Fig. 2b). Note that in the circular arrangement of Fig. 2b there is no telomeric (3’) or centromeric (5’) end of the Hox gene cluster and this may lead to the circular arrangement of the modules of the sea urchin larva. The circular gene arrangement is transient and in the course of gene restructuring, the closed contour returns to a linear configuration (see below). Furthermore, the closed gene fiber should not be confused with any circular DNA sequence observed in prokaryotic organisms like viruses or bakteria.

In order to keep up with the circular description it is useful to use a clockface notation of the polar coordinates with the 12 ‘hours’ representing 12 theoretical ontogenic modules arranged on a circle (Fig.3a). The same clockface notation is used for the closed Hox gene cluster in the larva stages. In this case the 12 ‘hours’ stand for the 12 Hox genes. (This is a formal simplification since the real sea urchin Hox genes are not 12 but 13). The discontinuity at the top (0/12) is artificial and can be easily handled. (See the Note at the end).

**Figure 3:**
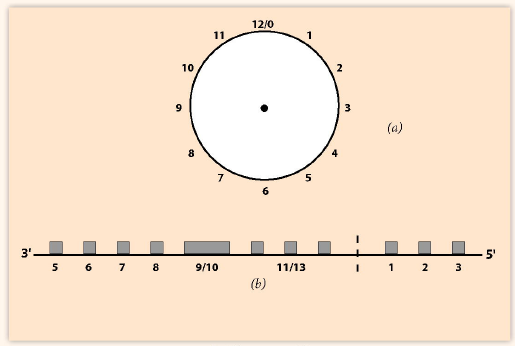
(a) Clockface representation of ‘hours’ 1-12. (b) Break and opening of the cluster at ‘hour’ (4) of Fig. 3a. Linear deployment of the cluster after the break at the dashed line.

## Symmetries of the sea urchin embryo and larva

At the first ontogenic stages of Echinoderms the anterior-posterior axis (A/P) is established before the larva stages [15]. This A/P axis runs from the mouth (the anterior end) through the adult coelomic compartments [16]. At the subsequent larva stages rotational symmetry is superimposed on the A/P axis [14].

For the different mathematical forms of Symmetry and particularly the applications in Echinoderms see the thought provoking book of H. Weyl [17]. The principal symmetries observed in both animals and plants are resulting from the two fundamental operations of *reflections* and *rotations* [17]: (a) a body (a geometric configuration) is *bilaterally symmetric* with respect to a plane P if it is carried into itself when reflected in P. (b) a body is *rotationally symmetric* around an axis L if it is carried into itself by a rotation around L [17]. The angle of rotation α may be α = (2π/n) where n = 1, 2, 3,…determines the *order* of rotation. For example, for n = 2 the body is symmetric for a rotation of 180**°**. For the needs of the present work it is not necessary to refer to the important case of three-dimensional bodies with left-right symmetry or asymmetry.

The echinoderm larva is rotationally symmetric for different angles of rotation. For example the pentamerous body structures of the sea urchin *H. purpurescens* with the five primary podia located on a circular arrangement around the axis is shown in Fig.2a [14]. An angular rotation (2π/5) around the axis translocates the podia: A to B, B to C, C to D, D to E and E back to A.

The circular structure of the sea urchin larva offers itself to a clockface representation (Fig.3a). At some point after the larva stage, the circular arrangement of the Hox cluster breaks and opens at the level of Hox4 [11,12] and the closed contour transforms to a linear gene sequence. The linear deployment of the genes is shown in Fig.3b. The 5’ side of this deployment (Hox1, Hox2, Hox3) is not satisfactory.

<u>Inversion of anterior genes:</u> Assume that podium D of Fig.2a extends in three distinct ‘hours’ (1, 2, 3) of the clockface representation. In the rotationally symmetric podium D these ‘hours’ are distributed on a local clockface (Fig.4a). The opening of this clockface at the dashed line leads to a linear arrangement between ‘hours’ (12/0) and (4) as shown in Fig.4b. Hox1, Hox2 and Hox3 are positioned on the side of (12/0) (gray squares) while the remaining ‘hours’ are not occupied (open squares) increasing thus the length of the Hox cluster. The resulting Hox gene ordering coincides with the unusual sea urchin Hox cluster of Fig.1b which violates the ‘normal’ Hox collinearity.

**Figure 4:**
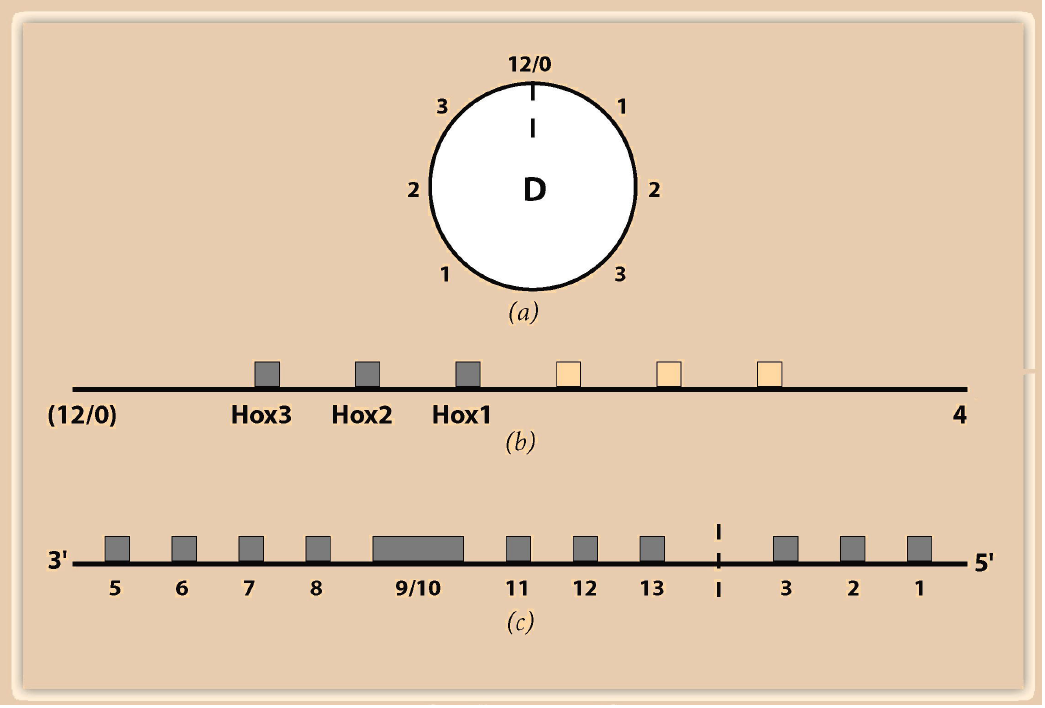
(a) Rotational symmetry of order 2 for podium D extending to ‘hours’ 1, 2, 3. (b) Break of the podium circle at the dashed line of Fig.4a and attachment of the line ends between ‘hours’ (12/0) and (4) of the larva clockface. Hox1, Hox2, Hox3 (full squares) are located at the (12/0) side. (c) Unusual linear deployment of the Hox cluster comparable to Fig.1b.

## Questions and Conclusions

In the present work, the particular case of the sea urchin is worked out as an example. The Hox clusters of the other clades can be treated similarly. The crucial evolutionary influences have not been considered. The effort is focused on the derivation of the unusual sequence of the echinoderm genes in the Hox cluster.

If the hypothesis of Hox gene congruence is confirmed, many questions will be raised to answer: what are the modes of echinoderm Hox gene expressions (in space and time)? Can any of the existing models (e.g. the biophysical model) describe the results? The existing data are for the moment rather fragmentary for a safe analysis.

### Note

The clockface representation was introduced some years ago in a quite different context, namely the ‘Polar Coordinate Model’ for epimorphic regeneration [18]. The twelve ‘hours’ 1-12 were used in the French, Bryant and Bryant model (FBB) to describe the supernumerary outgrowths developing after contralateral graftings e.g.in amphibian limbs. The FBB model was based on the <u>continuity</u> hypothesis of the shortest path of the intercalating clockface positional values. However, the Polar Coordinate Model failed to reproduce the outgrowths after 180° ipsilateral limb rotations. A complementary ‘Hierarchical Polar Coordinate Model’ could deal with the results after both contralateral and ipsilateral graftings [19]. This extended model was based on the hypothesis of continuity and congruence of the intercalating positional values.

